# Relative contribution of non-structural protein 1 in dengue pathogenesis

**DOI:** 10.1101/2020.02.01.929885

**Authors:** Pei Xuan Lee, Donald Heng Rong Ting, Clement Peng Hee Boey, Eunice Tze Xin Tan, Janice Zuo Hui Chia, Li Ching Ong, Yen Leong Chua, Chanditha Hapuarachchi, Lee Ching Ng, Sylvie Alonso

**Author notes:** Correspondence: 28 Medical Drive, Life Sciences Institute, Immunology programme, Singapore 117456.

## Abstract

Dengue is a major public health concern in the tropical and sub-tropical world with no effective treatment. The controversial live attenuated virus vaccine Dengvaxia has boosted the pursuit of sub-unit vaccine approaches, and the non-structural protein 1 (NS1) has recently emerged as a promising candidate. However, we found that NS1 immunization or passive transfer of NS1 antibodies failed to confer protection in symptomatic dengue mouse models using two non mouse-adapted DENV2 strains from the Cosmopolitan genotype that currently circulates in South-East Asia. Furthermore, exogenous administration of purified NS1 did not worsen *in vivo* vascular leakage in sub-lethally infected mice, thereby supporting that NS1 does not play a critical role in the pathogenesis of these DENV2 strains. Virus chimerization approaches indicated that the prME structural region, but not NS1, plays a critical role in driving *in vivo* fitness and virulence of the virus, through induction of key pro-inflammatory cytokines. This work highlights that the pathogenic role of NS1 is DENV strain-dependent, which warrants re-evaluation of NS1 as a universal dengue vaccine candidate.

## Introduction

Dengue virus (DENV) is a mosquito-borne virus responsible for an estimated 390 million annual infections in the tropical and sub-tropical world (Bhatt et al., 2013). The virus exists as four antigenically distinct serotypes (DENV1-4). Infection with DENV results in a wide spectrum of disease manifestations ranging from mild to life-threatening conditions, the latter being characterized by vascular leakage with or without hemorrhage development (World Health Organisation, 2009, Kyle and Harris, 2008). DENV serotype cross-reactive immunity has been proposed, and to a certain extent demonstrated, to represent a risk for the development of severe dengue. Specifically, pre-existing antibodies raised during a previous heterotypic DENV infection that recognize the structural components of the virion could enhance DENV uptake and facilitate its replication within FcγR-bearing cells, leading to increased disease severity, a phenomenon coined as antibody-dependent enhancement (ADE) (Halstead et al., 2002, Halstead, 2003). There is currently no effective therapeutics or vaccine against dengue, which mainly stems from our lack of understanding of dengue pathogenesis associated with limited availability of relevant symptomatic animal models (Yauch and Shresta, 2008, Chan et al., 2015). The only licensed vaccine Dengvaxia®, which consists of chimeric live attenuated tetravalent DENV, not only offers limited protective efficacy particularly towards DENV2 strains, but was also found to predispose immunologically dengue naïve individuals to an increased risk of developing severe disease (Capeding et al., 2014, Villar et al., 2015, Aguiar et al., 2016, Halstead, 2017). This latter observation had led to the suspension of dengue vaccination programmes in the Philippines (Fatima and Syed, 2018). Thus, safer and more effective second-generation vaccines are urgently needed. In the wake of Dengvaxia® setback and controversy, sub-unit vaccine candidates have regained some traction (Lam et al., 2016). While the envelope (E) protein of DENV has been the most popular sub-unit vaccine candidate, a number of studies have also proposed DENV non-structural protein 1 (NS1) as a potential candidate (Schlesinger et al., 1987, Henchal et al., 1988, Falgout et al., 1990, Beatty et al., 2015, Goncalves et al., 2015, Wan et al., 2014, Wan et al., 2017, Lai et al., 2017). NS1 is a glycosylated non-structural protein (46-55kDa) that homo-dimerizes after post-translational modification, becomes membrane-associated and participates in RNA replication within membrane-bound replication complexes. Soluble hexameric NS1 (sNS1) is made of three dimers and a central lipid cargo, and is secreted into the extracellular milieu (Gutsche et al., 2011, Muller and Young, 2013, Watterson et al., 2016). Cell surface-associated NS1 dimers and sNS1 hexamers are highly immunogenic (Muller and Young, 2013). sNS1 is detected in the blood at concentration of up to 50 μg/mL and typically follows viremia in dengue patients (Young et al., 2000, Libraty et al., 2002, Alcon et al., 2002), thus widely used as an early diagnostic marker (Peeling et al., 2010, Hang et al., 2009, Chuansumrit et al., 2011). Furthermore, recent studies have reported a role for sNS1 in dengue pathogenesis, specifically in causing vascular leakage, a key clinical manifestation of severe dengue (Beatty et al., 2015, Modhiran et al., 2015). As such, immunity raised against NS1, either through direct immunization with purified sNS1 or via passive transfer of anti-NS1 immune serum, was found to afford protection against lethal DENV infection in a mouse model (Beatty et al., 2015). These findings thus support that NS1 represents a promising sub-unit vaccine candidate and a safer alternative to avoid the risk of ADE, which is a major roadblock in the development of vaccines that rely on protective immunity against the structural components of DENV. However, the role of NS1 in dengue pathogenesis and the *in vivo* protective efficacy of NS1 immunity had been demonstrated in mouse models that either produce mild symptoms such as local skin hemorrhage and prolonged bleeding time (Wan et al., 2014, Wan et al., 2017, Lai et al., 2017) or employ mouse-adapted DENV2 strains (Schlesinger et al., 1987, Henchal et al., 1988, Falgout et al., 1990, Costa et al., 2006, Beatty et al., 2015).

Here, the role of NS1 in dengue pathogenesis as well as the protective potential of NS1 immunity were investigated in symptomatic dengue mouse models established with a non mouse-adapted DENV2 strain, namely D2Y98P (Tan et al., 2010, Ng et al., 2014, Martinez Gomez et al., 2016). In these models, neutralization of circulating sNS1 through antibody transfer or active NS1 immunization did not confer protection against D2Y98P-induced disease, thus suggesting a limited role of sNS1 in the pathogenesis of D2Y98P. Similar observations were made with a DENV2 clinical isolate that currently circulates in Singapore and Malaysia. Instead, we showed that the prME region drives the *in vivo* virulence of D2Y98P virus, through induction of key pro-inflammatory cytokines.

## Results

### No protective efficacy of NS1 immunity in A129 and AG129 ADE mouse models

The protective efficacy of NS1 was evaluated in symptomatic mouse models of dengue that have been established with the non-mouse-adapted DENV2 D2Y98P strain (Tan et al., 2010, Ng et al., 2014, Martinez Gomez et al., 2016). D2Y98P derives from a 1998 clinical isolate from the DENV2 Cosmopolitan genotype and has been exclusively passaged for approximately 20 rounds in C6/36 mosquito cells (Tan et al., 2010). E protein nucleotide sequence or the entire coding sequence of D2Y98P genome was compared to that of a DENV2 prototype strain – NGC strain (Anez et al., 2016), and with six other DENV2 Singapore isolates that were collected between 2007-2010 (Lee et al., 2012). The results indicated 94% identity with NGC and 98% identity with the six Singapore isolates, thus supporting that D2Y98P can be considered as a representative of DENV2 strains that circulate in Singapore (Table S1).

Adult A129 mice (deficient in Type I interferon (IFN) pathway) were administered thrice with commercially available purified hexameric NS1 from DENV2 strain 16681 according to a previously published immunization protocol (Beatty et al., 2015). High titers of NS1-specific IgG antibodies were measured in the serum of the immunized mice, with majority of the IgG1 subclass (Fig. S1). A control group consisting of mice administered with ovalbumin (OVA) according to the same immunization regimen was included. OVA- and NS1-immunized mice were then administered with DENV1-immune serum one day prior to infection with D2Y98P strain in order to produce lethal infection in an ADE setting (Fig. 1A).

**Fig. 1.**
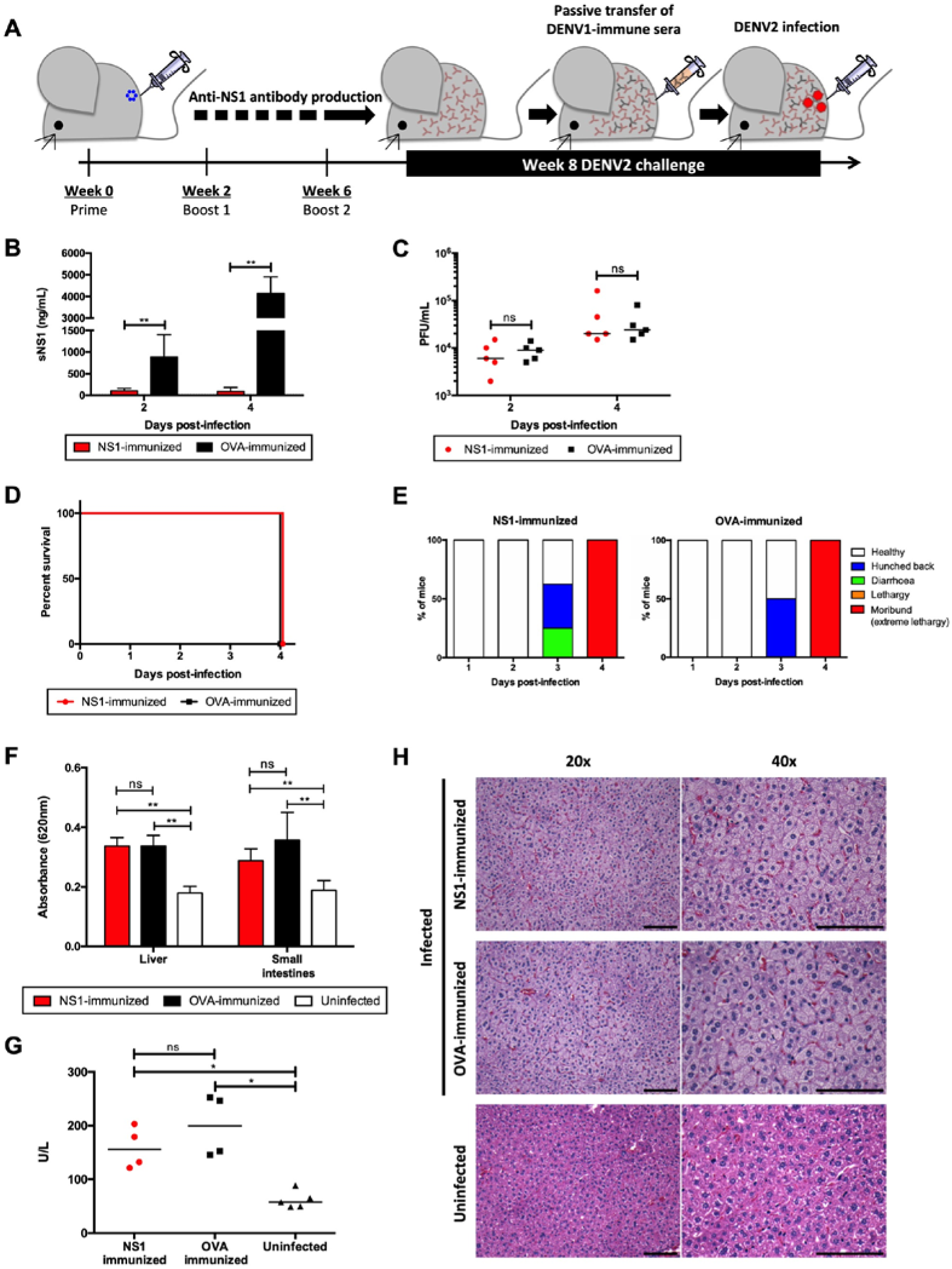
NS1 immunization in A129 ADE infection model. (A) A129 mice were immunized (ip.) thrice with 20 μg NS1(16681) or OVA adjuvanted with MPLA and Addavax. Eight weeks post-immunization, mice were passively transferred with DENV1-immune serum one day before iv challenge with 10^6^ PFU DENV2 (D2Y98P). (B) Circulating sNS1 levels and (C) viremia titers were measured (n = 5) at day 2 and 4 p.i. (D, E) Mice (n = 8) were monitored daily upon challenge. The Kaplan-Meier survival curve and clinical scores are shown. (F) Vascular leakage was quantified by Evans blue dye extravasation assay in liver and small intestines at day 4 p.i. (n = 5). (G) Systemic AST levels were measured (n = 4-5) and (H) histological analysis of liver was performed at day 4 p.i. (n = 4-5). Images were taken at 20x and 40x magnification. Representative sections are shown. (Scale bar – 10 μm). Data were analyzed by non-parametric Mann Whitney test. *\*p*<0.05; *\*\*p*<0.01; ns: not significant.

NS1 (16681) and NS1 (D2Y98P) share 97% amino acid identity and ability of the immune serum raised against NS1(16681) to bind to and neutralize NS1(D2Y98P) was verified by ELISA and sandwich ELISA, respectively (Fig. S1B and S1D). Consistently, significant reduction in circulating sNS1 was observed in NS1-immunized mice compared to OVA-immunized animals after viral challenge (Fig. 1B), indicating that NS1(16681) immunization successfully neutralized circulating sNS1 produced during infection with D2Y98P virus. However, comparable viremia titers were measured between NS1-immunized mice and OVA controls (Fig. 1C). Furthermore, NS1- and OVA-immunized mice displayed similar disease manifestations and progression, including hunched back, diarrhoea, lethargy, and eventually all the animals from both groups were moribund by day 4 post-infection (p.i.) (Fig. 1D and E). Vascular leakage in the liver and small intestines from NS1-immunized animals was as extensive as that measured in the OVA control group (Fig. 1F). Systemic level of the liver enzyme AST was also comparable between NS1- and OVA-immunized groups (Fig. 1G) and massive cytoplasmic vacuolation was apparent in the hepatocytes from both infected groups (Fig. 1H), indicating that reduction in sNS1 levels brought about by NS1 immunization did not protect the mice from DENV-induced severe liver damage (Martinez Gomez et al., 2016).

Therefore, despite effective neutralization of circulating sNS1, active immunization with purified NS1 failed to provide protection against lethal DENV2 challenge in an A129 ADE model.

The protective potential of NS1 immunity was further explored in another lethal ADE model using AG129 mice (deficient in Type I & II IFN pathways) – a model in which mice born to DENV1-immune dams exhibit extensive vascular leakage upon DENV2 infection (Ng et al., 2014). Since AG129 mice lack a functional IFNγ signalling pathway, direct immunization with purified NS1 may result in sub-optimal immune responses that may affect the protective potential of NS1 immunity (van den Broek et al., 1995). To address this possibility, NS1 immune serum was instead generated in A129 mice according to the same immunization protocol described above. One day prior to DENV2 infection, AG129 mice born to DENV1-immune dams were then passively transferred with the NS1 immune serum (Fig. 2A), which led to effective neutralization of circulating sNS1 after D2Y98P challenge (Fig. 2B). However, comparable viremia titers (Fig. 2C), survival rate (Fig. 2D), clinical symptoms (Fig. 2E) and vascular leakage (Fig. 2F) were observed between mice administered with NS1 immune serum and the control group transferred with naïve serum.

**Fig. 2.**
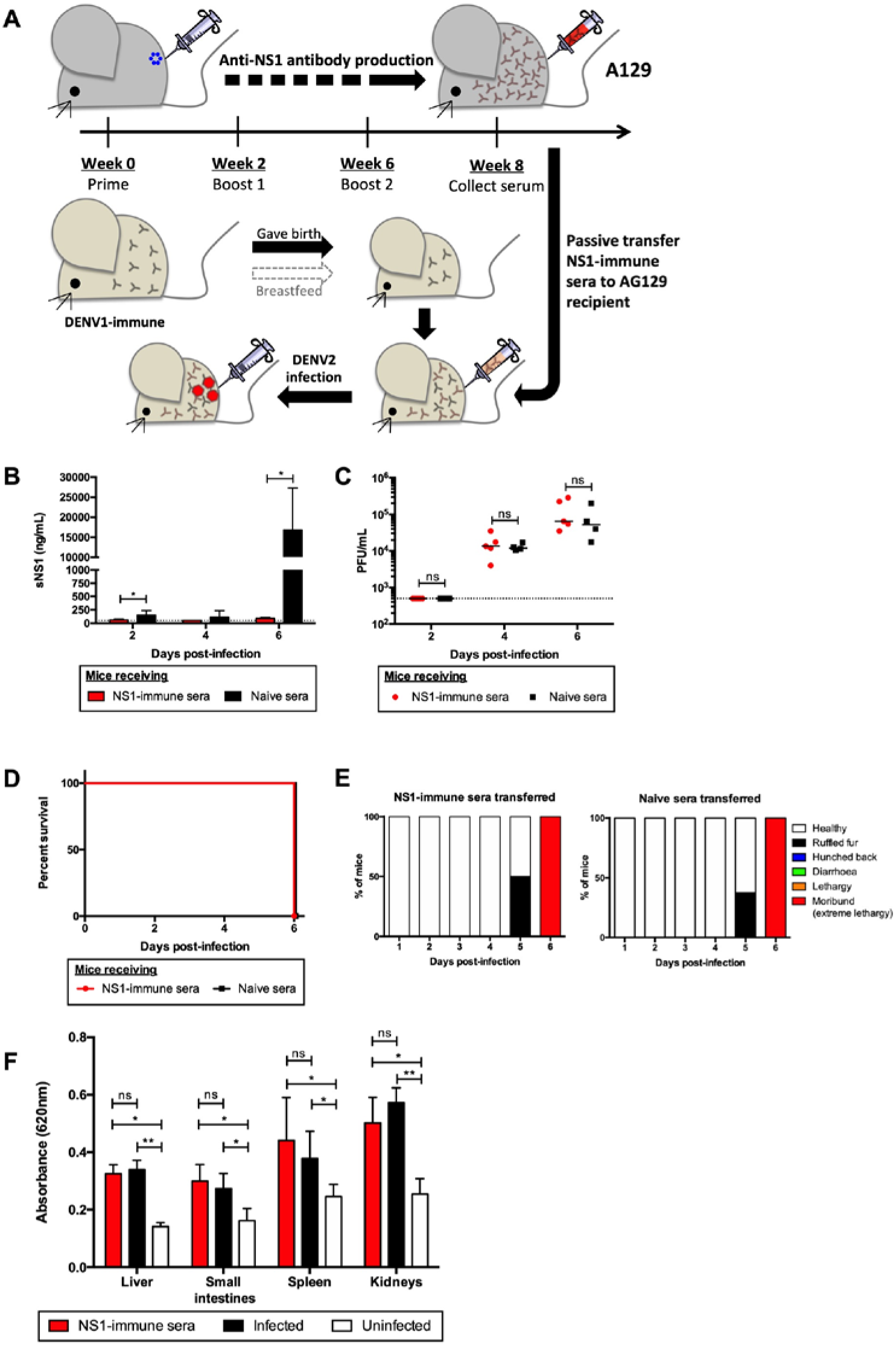
Passive NS1 immunity in AG129 ADE infection model. (A) AG129 mice born to DENV1-immune mothers were passively transferred with NS1 immune serum collected from NS1(16681)-immunized A129 mice (Fig. 1A) or with naïve serum. One day after transfer, the mice were sc challenged with 10^3^ PFU D2Y98P. (B) Systemic sNS1 levels and (C) viremia titers were measured at day 2, 4 and 6 p.i. (n = 4-5). (D, E) Mice (n = 8) were monitored daily upon challenge. The Kaplan-Meier survival curve and clinical score were shown. (F) Vascular leakage was assessed in indicated tissues at day 6 p.i. (n = 4-5). Data were analyzed by non-parametric Mann Whitney test. *\*p*<0.05; *\*\*p*<0.01; ns: not significant.

Together, our data indicate that neither active NS1 immunization nor passive transfer of NS1 immune serum conferred significant protection in both A129 and AG129 ADE models, despite effective neutralization of circulating sNS1. These observations thus imply that in these ADE infection models, sNS1 does not play a critical role in dengue pathogenesis.

### Lack of protective NS1 immunity is independent of mouse background

Previous studies that reported NS1-mediated protection were conducted in mouse strains such as IFNAR^-/-^ (Beatty et al., 2015) and STAT1^-/-^ (Wan et al., 2017, Lai et al., 2017), both being in the C57BL/6 background. We thus questioned whether the discrepancy between our observations and these earlier studies could be explained by a difference in mouse background. Hence, IFNAR^-/-^ mice of C56BL/6 background were immunized with NS1 or OVA according to the same immunization regimen as above, followed by lethal challenge with D2Y98P (Fig. 3A). NS1-immunized IFNAR^-/-^ mice had significantly reduced sNS1 levels after D2Y98P challenge compared to OVA-immunized controls, indicating effective neutralization of sNS1 in circulation (Fig. 3B). However, comparable viremia titers (Fig. 3C), survival rates (Fig. 3D), weight loss profiles (Fig. 3E) and vascular leakage (Fig. 3F) were measured between NS1-immunized and OVA control groups. These data thus indicate that the lack of protection mediated by NS1-immunity is likely independent of the mouse background.

**Fig. 3.**
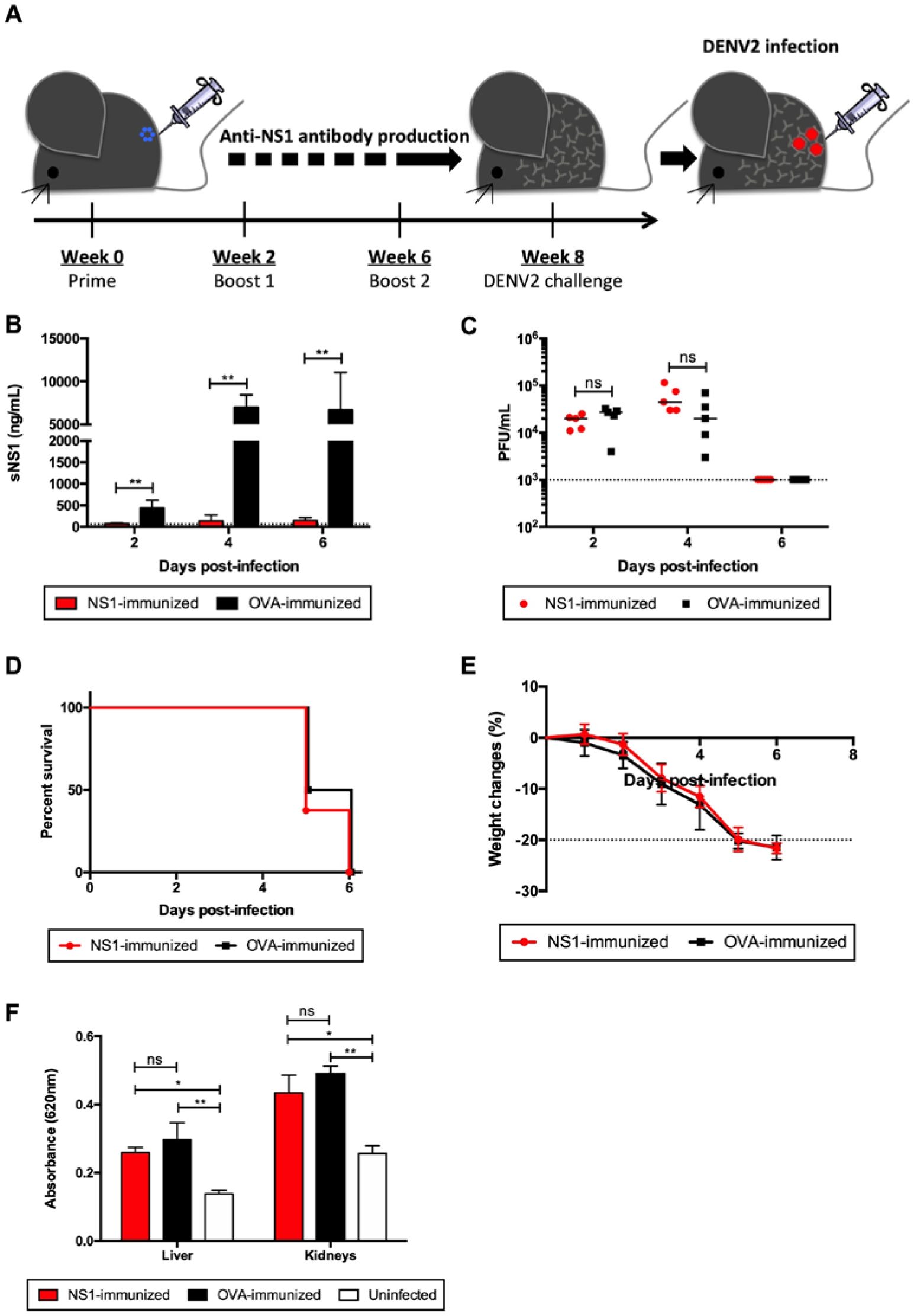
NS1 immunization in IFNAR^-/-^ primary infection model. (A) IFNAR^-/-^ mice were immunized with NS1(16681) or OVA as described in the legend of Fig. 1. Eight weeks post-immunization, mice were sc challenged with 10^5^ PFU D2Y98P. (B) Systemic sNS1 levels and (C) viremia titers were measured at day 2, 4 and 6 p.i. (n = 5). (D, E) Mice (n = 6-8) were monitored daily upon challenge. The Kaplan-Meier survival curve and weight loss profile are shown. (F) Vascular leakage was assessed in liver and kidneys at day 4 p.i. (n = 4-5). Data were analyzed by non-parametric Mann Whitney test. *\*p*<0.05; *\*\*p*<0.01; ns: not significant.

### Lack of NS1-mediated protective efficacy was also observed with another circulating DENV2 Singapore clinical isolate

While D2Y98P is not mouse-adapted, the virus had been passaged in C6/36 cell line for several rounds, during which some mutations in the viral genome may have modified the virus fitness. In particular, the presence of a phenylalanine (F) at position 52 in NS4B was found to play an important role in D2Y98P *in vivo* and *in vitro* fitness (Grant et al., 2011, Tan et al., 2010). As none of the other DENV2 strains whose genome sequence is available in the public database, harbors an F at this position, we proposed that this particular amino acid was likely acquired during *in vitro* passages. To examine whether the findings made with D2Y98P can be extended to other DENV2 strains, we have used another DENV2 clinical isolate, EHIE2862Y15, which has been passaged in C6/36 cell line for no more than four rounds. This virus belongs to the cosmopolitan genotype clade Ib that has been circulating in Singapore and Malaysia since 2013 and has been associated with high fatality rate (Ng et al., 2015). EHIE2862Y15 shares 98.6% nucleotide identity and 99.6% amino acid identity with D2Y98P (Table S2). Interestingly, although EHIE2862Y15 does not harbor an F at position 52 in NS4B, infection with EHIE2862Y15 gave rise to symptomatic infection in AG129 mice, with a disease kinetic profile similar to that of D2Y98P (Fig. S2).

To evaluate the role of NS1 in the context of EHIE2862Y15 infection, IFNAR^-/-^ mice were immunized with purified NS1 followed by lethal viral challenge. Despite the effective neutralization of circulating sNS1 (Fig. 4A), mice were not protected from lethal EHIE2862Y15 challenge, as evidenced by comparable viremia titers (Fig. 4B), survival rates (Fig. 4C) and weight loss profile (Fig. 4D). Furthermore, significant vascular leakage was measured in the liver and kidneys from NS1- and OVA-immunized mice compared to uninfected controls, although the extent of vascular leakage in the liver of NS1-immunized mice was slightly but significantly lower than in OVA-immunized animals (Fig. 4E). Overall, these data support that the lack of NS1 immunity-mediated protection is not restricted to D2Y98P strain, but also applies to a clinically relevant DENV2 isolate that is circulating in Singapore.

**Fig. 4.**
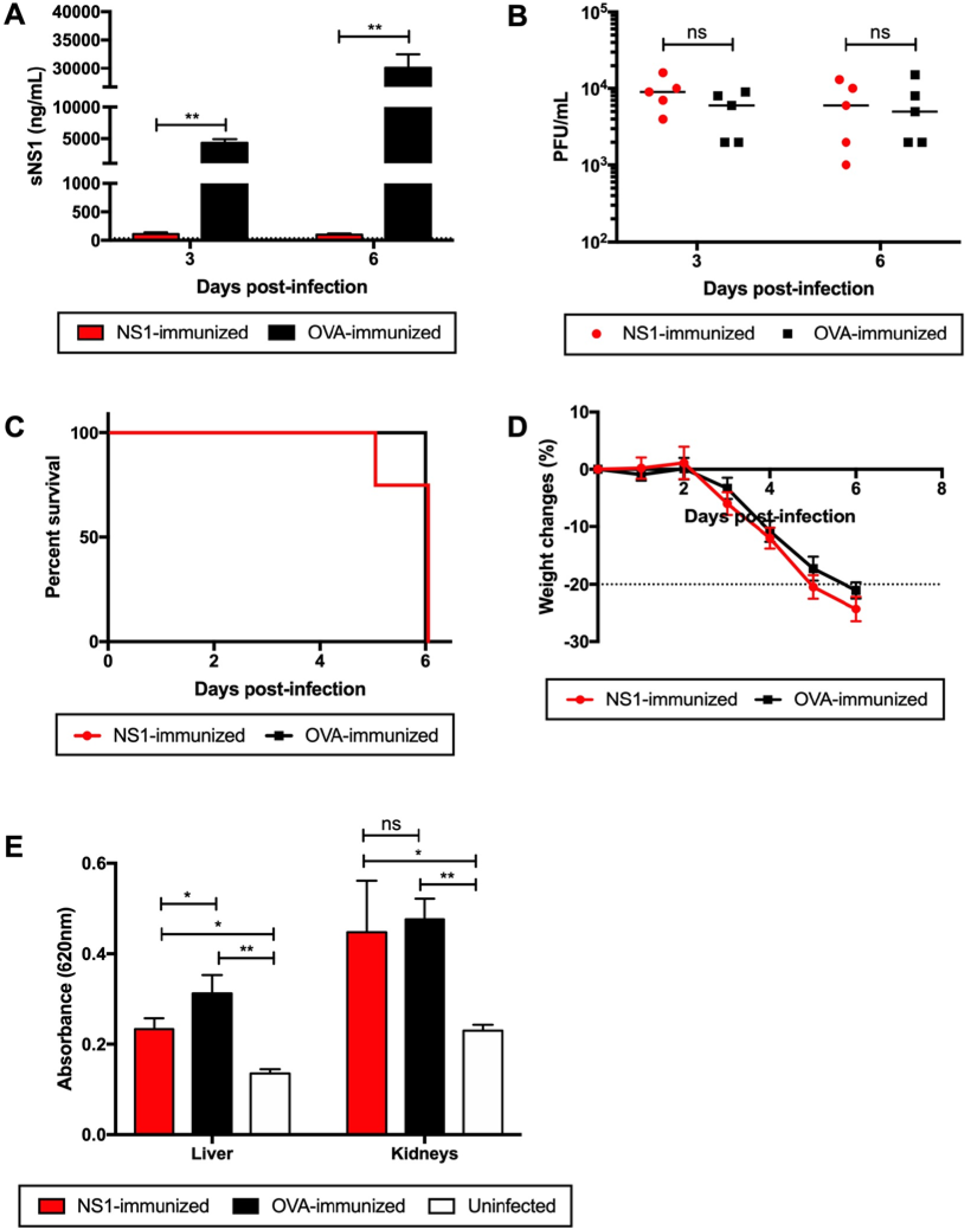
Protective efficacy of NS1 immunity against a low-passaged DENV2 clinical isolate. IFNAR^-/-^ mice were immunized with NS1(16681) or OVA as described in the legend of Fig 1. Eight weeks post-immunization, mice were sc challenged with 10^5^ PFU EHIE2862Y15. (A) Systemic sNS1 levels and (B) viremia titers were measured at day 3 and 6 p.i. (n = 5). (C, D) Mice (n = 8) were monitored daily upon challenge. The Kaplan-Meier survival curve and weight loss profile are shown. (E) Vascular leakage was assessed in the liver and kidneys at day 4 p.i. (n = 4-5). Data were analyzed by non-parametric Mann Whitney test. *\*p*<0.05; *\*\*p*<0.01; ns: not significant.

### Exogenous administration of sNS1 did not worsen dengue disease severity

To assess the pathogenic role of sNS1 during D2Y98P infection, we tested whether the exogenous administration of purified sNS1(D2Y98P) could aggravate disease severity in AG129 mice infected with a sub-lethal dose of D2Y98P (Fig. 5A). Administration of purified sNS1(D2Y98P) at day 2 p.i. led to significantly higher sNS1 level measured at day 4 p.i. compared to mice administered with OVA (Fig. 5B). No significant vascular leakage however was detected in any of the infected mice from both groups at this time point (Fig. 5C). Vascular leakage was also measured at day 6 p.i., when comparable levels of sNS1 were measured in both groups (Fig. 5B). It is likely that at this later time point, the sNS1 level mainly results from virus replication, and that exogenously added sNS1 had been cleared from the circulation and/or deposited onto the endothelial layer and internalized by endothelial cells. While no significant vascular leakage was observed in the small intestine and spleen from both infected groups, liver and kidneys did display significant vascular leak compared to uninfected controls (Fig. 5D). However, the extent of vascular leakage was not greater in mice administered with exogenous sNS1 compared to the OVA control group (Fig. 5D).

**Fig. 5.**
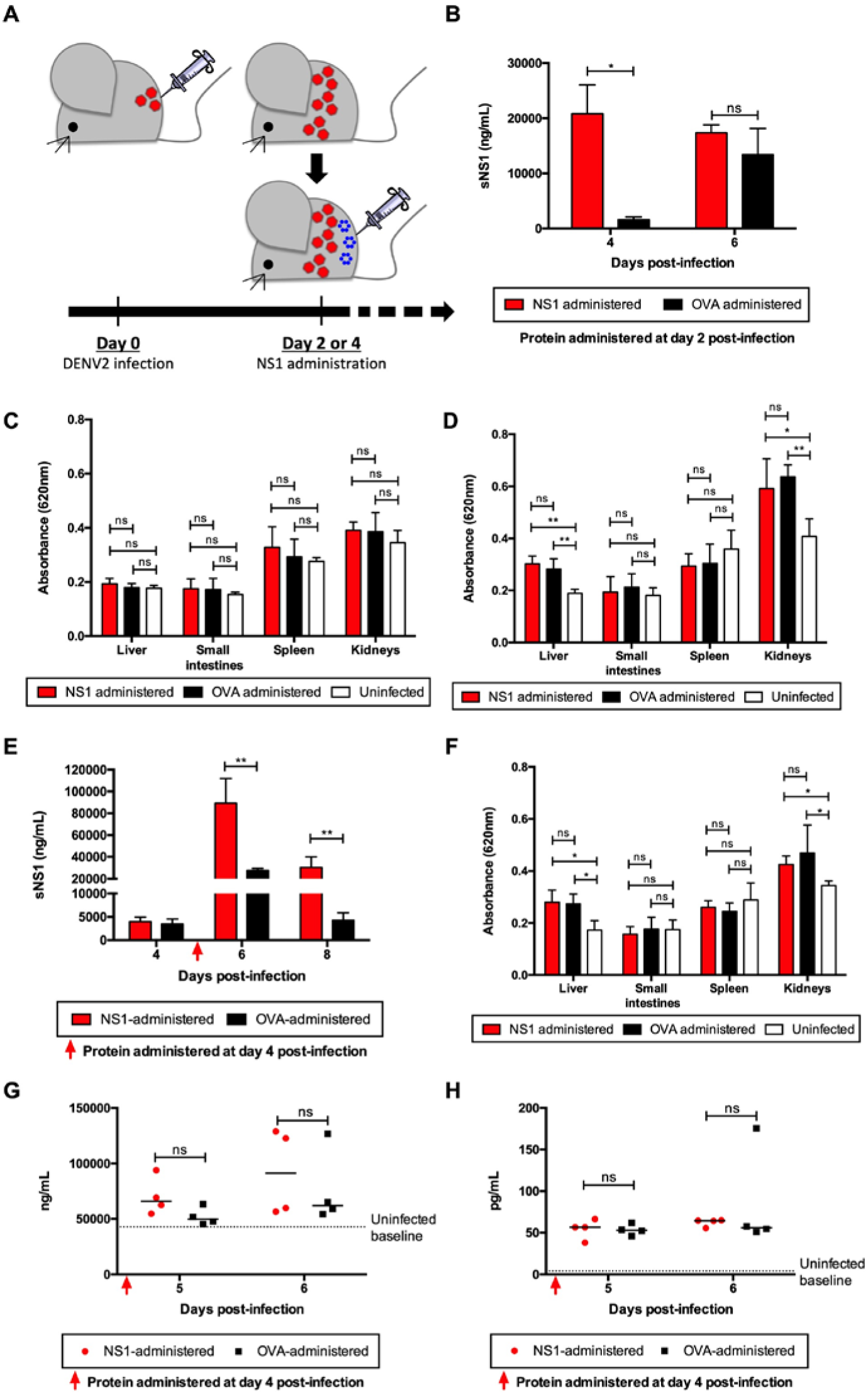
Administration of NS1(D2Y98P) to AG129 mice infected with a sub-lethal dose of D2Y98P. (A) AG129 mice were sc challenged with 10^3^ PFU D2Y98P. At (B-D) day 2 p.i. or (E-H) day 4 p.i., infected mice were iv administered with 10mg/kg NS1(D2Y98P) or OVA. (B, E) Systemic levels of sNS1 were measured at various time points p.i. (n = 4-5). Vascular leakage was evaluated in various tissues at (C) day 4 or (D, F) day 6 p.i. (n = 4-5). (G) Plasma heparan sulphate (n = 4) and (H) TNF-α levels (n = 4) were measured at day 4 p.i. Data were analyzed by non-parametric Mann Whitney test. *\*p*<0.05; *\*\*p*<0.01; ns: not significant.

Since vascular leakage is more prominent at day 6 p.i. in this sub-lethal AG129 model, we reasoned that delaying the exogenous administration of sNS1 from day 2 p.i. to day 4 p.i. may have a greater impact on vascular leakage. Significantly higher circulating sNS1 levels were measured at day 6 p.i. in mice administered with purified sNS1 (Fig. 5E). However, these mice failed to exhibit greater vascular leakage than the OVA control group (Fig. 5F). Viremia titers were also not enhanced with sNS1 administration (Fig. S3).

Previous studies have reported the ability of sNS1 to activate toll-like receptor-4 (TLR4) on immune cells leading to the production of vasoactive cytokines TNF-α (Modhiran et al., 2015) and causing endothelial glycocalyx disruption (Puerta-Guardo et al., 2016). However, D2Y98P-infected mice administered with exogenous sNS1 did not display significantly higher levels of TNF-α compared to OVA controls (Fig. 5G). Likewise, the levels of heparan sulphate in circulation, indicative of glycocalyx degradation, were not enhanced with NS1 administration (Fig. 5H). These findings are consistent with the observation that exogenous administration of sNS1 did not increase vascular leakage in D2Y98P-infected mice.

### *In vivo* virulence of D2Y98P is driven by structural prME region

The data described above supported that NS1 does not play a critical role in the pathogenesis and *in vivo* fitness of the two DENV2 strains employed, and that other viral determinants are involved. The *in vivo* fitness of DENV was reported to be linked to its organ and cell tropism, which is mainly mediated by the prME structural region that harbors the receptor-binding site (Modis et al., 2004, Prestwood et al., 2008). To explore the mechanisms involved in D2Y98P fitness and virulence, we generated a chimeric virus where the prME region from D2Y98P was swapped with that of a DENV1 strain – 05K3903DK1, which typically gives rise to asymptomatic transient viremia in AG129 mice (Fig. 6G). We also engineered another chimeric virus where NS1 from D2Y98P was replaced by that of DENV1 (Fig. 6A and B).

**Fig. 6.**
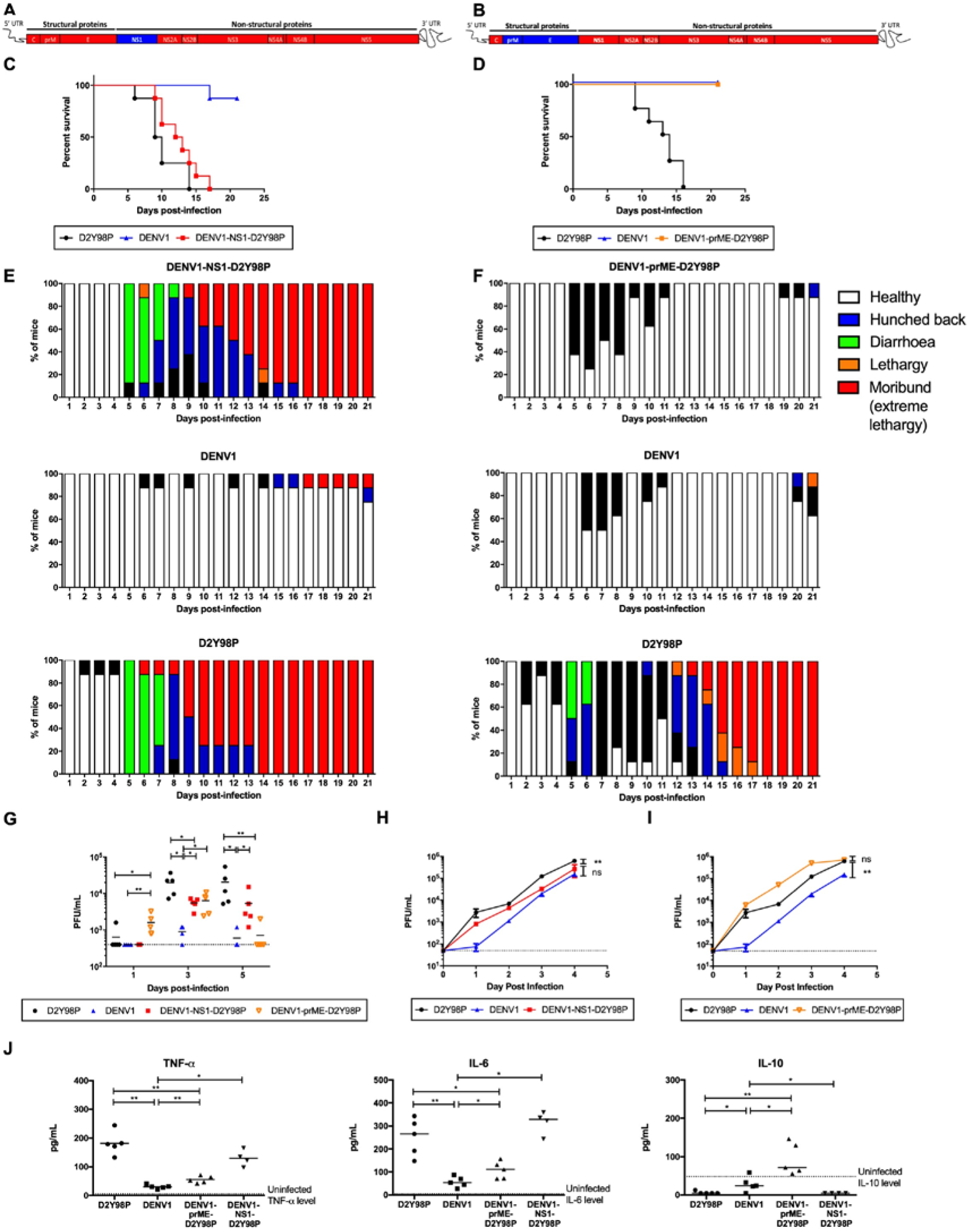
*In vivo* and *in vitro* characterization of DENV1-NS1-D2Y98P and DENV1-prME-D2Y98P chimeric viruses. (A) DENV1-NS1-D2Y98P or (B) DENV1-prME-D2Y98P chimeric strains consist of (A) NS1 or (B) prME coding region from DENV1 strain (blue) cloned into the backbone of D2Y98P strain (red). AG129 mice were sc infected with 10^6^ PFU of D2Y98P, DENV1, DENV1-NS1-D2Y98P or DENV1-prME-D2Y98P. Mice (n = 8) were monitored daily upon challenge. (C, D) The Kaplan-Meier survival curves and (E, F) clinical scores are shown. (G) Viremia titers were measured at day 1, 3 and 5 p.i. (n = 4). (H, I) *In vitro* virus titer in Vero cells infected with respective viruses. The viral kinetic curves were compared and analyzed by linear regression, which takes into consideration the slope and Y-intercept. (J) TNF-α, IL-6 and IL-10 systemic levels were measured at day 5 p.i. (n = 4-5). Data between WT and chimeric viruses were compared and analyzed by non-parametric Mann-Whitney test. \**p*<0.05; \*\**p*<0.01; ns: not significant.

Mice infected with DENV1-NS1-D2Y98P displayed a survival profile, clinical manifestations and disease progression that were comparable to those seen with WT D2Y98P virus (Fig. 6C, E). On the other hand, mice infected with DENV1-prME-D2Y98P remained asymptomatic throughout the experiment, similar to WT DENV1 (Fig. 6D, F). Interestingly, the peak viremia titer (day 3 p.i.) in mice infected with DENV1-prME-D2Y98P was similar to that measured in mice infected with DENV1-NS1-D2Y98P (Fig. 6G), although the infection outcome induced by both viruses differed drastically. Together, these observations support that prME, but not NS1, drives D2Y98P virulence in mice. Swapping either NS1 or prME from DENV1 into D2Y98P did impact significantly the viremia titers (Fig 6G), suggesting that these proteins play a role in viral replication. The role of NS1 in DENV replication has indeed been well established (Fan et al., 2014). Furthermore, a previous work has also reported the existence of specific interactions between NS1 and prM/E that facilitate membrane budding or conformational changes in the structural proteins necessary for formation of nucleocapsids (Scaturro et al., 2015). This suggests that some compatibility between both proteins is required for optimal viral replication. Consistently, when comparing the kinetic profile in Vero cells of D2Y98P with that of DENV1-NS1-D2Y98P, a mild but significant defect could be observed with the chimeric strain (Fig. 6H). In contrast, the *in vitro* kinetic profile of DENV1-prME-D2Y98P was super-imposable to that of WT D2Y98P (Fig. 6I). This latter observation thus implied that the lower viremia titers measured in mice infected with DENV1-prME-D2Y98P cannot be explained by an intrinsic defect in viral replication.

To further understand the basis of the prME-driven virulence of D2Y98P, we examined the cytokine profile in mice infected with the chimeric and WT strains (Fig 6J and S4). Comparable high levels of key pro-inflammatory cytokines TNF-α and IL-6 were measured in mice infected with DENV1-NS1-D2Y98P and WT D2Y98P (Fig. 6J). On the other hand, mice infected with DENV1-prME-D2Y98P displayed significantly reduced levels of these pro-inflammatory cytokines compared to D2Y98P-infected mice, although higher than the levels measured in DENV1-infected animals (Fig. 6J). Furthermore, DENV1-prME-D2Y98P was unable to suppress the production of anti-inflammatory IL-10 (Fig. 6J).

Therefore, our data suggest that the *in vivo* virulence of D2Y98P virus is mainly driven by its structural component – prME – through its ability to induce the production of key pro-inflammatory cytokines.

## Discussion

The mechanisms that drive the fitness and virulence of DENV remain largely unknown and are likely to be multifactorial. They include the ability to infect and replicate effectively in various host cells as well as the ability to evade or counteract the host defences (Mukhopadhyay et al., 2005, Hsieh et al., 2014, Hsieh et al., 2011, Munoz-Jordan et al., 2003, Munoz-Jordan et al., 2005).

Beyond its role in DENV intracellular replication, recent literature has reported a critical role of NS1 in dengue pathogenesis by interacting with the endothelium and inducing vascular leakage, a clinical feature of severe dengue (Beatty et al., 2015, Modhiran et al., 2015). Consequently, NS1 immunity was found to protect against dengue disease (Beatty et al., 2015, Costa et al., 2006, Lai et al., 2017, Wan et al., 2014, Wan et al., 2017). These *in vivo* studies were conducted with DENV clinical isolates or mouse-adapted strains which either required very high dosage to induce lethality (DENV2-454009A, D220, DENV2-3295) or induced only mild clinical symptoms (DENV2-16681, DENV1-8700828, DENV3-8700829, DENV4-59201818, DENV2-RJ) (Orozco et al., 2012, Beatty et al., 2015, Costa et al., 2006, Lai et al., 2017, Wan et al., 2014, Wan et al., 2017, Chan et al., 2019), thus implying that the *in vivo* fitness of these DENV strains is rather poor.

Here, we have used a non mouse-adapted DENV2 strain (D2Y98P) that is highly virulent in mice (Tan et al., 2010, Ng et al., 2014, Martinez Gomez et al., 2016). In that context, and using several symptomatic models in different mouse backgrounds, we did not observe a role for circulating sNS1 in dengue pathogenesis. Neither could we see a protective role for NS1 immunity. Similar observations were made with another DENV2 clinical isolate that currently circulates in Singapore and Malaysia, and that is equally virulent in mice.

Consistently, the role of NS1 in human dengue disease has remained controversial. A hospital-based study in dengue patients reported a positive association between circulating sNS1 levels and dengue disease severity, and proposed that levels greater than 600 ng/mL are predictive of severe dengue (Libraty et al., 2002). However, a closer examination of the data revealed that the correlation was not absolute, where a proportion of patients with self-limiting dengue fever displayed levels of sNS1 above 600 ng/mL. Likewise, a subset of patients who developed severe dengue had sNS1 levels below the detectable range or considerably lower than 600 ng/mL (Libraty et al., 2002). Furthermore, a more recent prospective study conducted in Vietnam found no association between sNS1 level and severe dengue (Fox et al., 2011). As these patients were likely to be infected with DENV from different serotypes and strains, the authors suggested that the correlation between sNS1 levels and disease severity may be strain-dependent. A separate prospective study indeed reported that the level of circulating sNS1 produced in infected individuals differs greatly across DENV serotypes and even among strains of the same DENV2 serotype (Duyen et al., 2011). Therefore, data from dengue patient cohorts do not seem to support a strong correlation between the level of circulating sNS1 and disease severity.

Currently, little is known about the viral determinants that influence DENV virulence in humans. A number of studies have reported that some lineages seem to be more virulent than others, based on their ability to replicate in their host and give high viremia titers (Rico-Hesse et al., 1997, Leitmeyer et al., 1999, Armstrong and Rico-Hesse, 2001, Cologna et al., 2005). DENV strains with greater replicative ability in their host are thought to spread more rapidly and successfully than those with a lower fitness (Guzman and Harris, 2015, Cologna et al., 2005). As a non-structural protein that plays a critical role in viral replication, NS1 has been naturally proposed to be a key determinant in driving the fitness of DENV, and studies have shown that single amino acid substitutions within NS1 could affect viral production and consequently disease severity (Rodriguez-Roche et al., 2005, Rodriguez-Roche et al., 2011, Chan et al., 2019). Other mechanisms have also been proposed and include the ability to evade, deceive or interfere with the host defences, through production of sub-genomic viral RNA (Finol and Ooi, 2019), immature viral particles (Rodenhuis-Zybert et al., 2010), or specific interactions between viral and host proteins (Munoz-Jordan et al., 2003, Munoz-Jordan et al., 2005).

Our data support that prME drives the *in vivo* virulence of D2Y98P. As prM and E form the outer protein shell of the virion, these proteins have been found to play a critical role in receptor binding and entry into the host cell, thereby driving virus tropism (Mukhopadhyay et al., 2005, Hsieh et al., 2014, Hsieh et al., 2011). Specifically, the two N-linked glycans on E protein at positions 67 and 153 have been shown to be crucial for binding to DC-SIGN and mannose receptors (Pokidysheva et al., 2006, Tassaneetrithep et al., 2003, Miller et al., 2008). Although both N-glycosylation sites are conserved among DENV strains, the sugar composition and branching structure present at each site differ among serotypes and even among strains within the same serotype (Yap et al., 2017). The sugar composition at the surface of the viral particles is likely to impact binding efficacy of the virus to its lectin-type host receptors, thereby influencing cell and organ tropism. In addition to receptor binding, glycans on E protein may also play an important role in shaping the host immune responses, through modulation of the lectin pathway of complement activation, and induction of pro-inflammatory cytokines through C-type lectin domain family 5 member A (CLEC5A) receptor activation (Chen et al., 2008). Using a chimerization approach, we showed that swapping prME region from DENV1 into D2Y98P virus did not modify the *in vitro* viral replication rate but significantly impaired the induction of pro-inflammatory cytokines *in vivo*, which resulted in asymptomatic disease. We have indeed previously shown that pro-inflammatory cytokines, in particular TNFα, play a critical role in D2Y98P-induced disease, whereby administration of anti-TNFα antibodies partially or completely protected mice from succumbing to the infection (Ng et al., 2014, Martinez Gomez et al., 2016). Whether the glycan structures on (DENV1)E protein differ from those found on (D2Y98P)E and are responsible for the phenotype observed remains to be determined experimentally and is beyond the scope of this study. It is however interesting to note that the prME region from DENV2 Singapore clinical isolate – EHIE2862Y15, shares 100% amino acid identity with prME of D2Y98P.

In conclusion, our work supports that the pathogenic role of sNS1 and consequently the protective efficacy of NS1 immunity are not seen in the context of infection with two DENV2 strains that belong to the cosmopolitan genotype, and whose virulence relies instead on their structural components. Hence, we propose that the pathogenic role of sNS1 is likely to be DENV strain-dependent. Given the highly diverse and vast number of DENV strains circulating in the population (Kyle and Harris, 2008, Lee et al., 2012, Hapuarachchi et al., 2016), disease pathology in infected individuals could be driven by different key viral determinants with sNS1 playing a major or minor role. Hence, it is critical to re-evaluate the protective potential of NS1 vaccination against a variety of circulating DENV strains in order to avoid repeating the Dengvaxia scenario.

## Materials & Methods

### Ethics statement

All the animal experiments were carried in accordance with the guidelines of the National Advisory Committee for Laboratory Animal Research (NACLAR). Animal facilities are licensed by the regulatory body Agri-Food and Veterinary Authority of Singapore (AVA). The described animal experiments were approved by the Institutional Animal Care and Use Committee (IACUC) from National University of Singapore (NUS) under the protocol number R14-0992 and R16-0422.

### Cell lines and viruses

C6/36 *Aedes albopictus* cell line (American Type Culture Collection (ATCC) #CRL-1660) was maintained in Leibotvitz’s L-15 medium (GIBCO) supplemented with 10% fetal bovine serum (FBS) (GIBCO) at 28 °C. Baby hamster kidney-21 (BHK-21) (ATCC #CCL-10) cell line was maintained in RPMI-1640 medium (GIBCO) supplemented with 10% FBS and cultured at 37 °C with 5% CO_2_. African green monkey kidney epithelial (Vero) (ATCC #CCL-81) cell line was maintained in DMEM medium (GIBCO) supplemented with 10% FBS and cultured at 37 °C with 5% CO_2_. Human dermal blood microvascular endothelial cells (HMVEC) (#CC-2811) (Lonza) were maintained in EGM™-2MV BulletKit™ medium (Lonza) at 37 °C with 5% CO_2_. DENV1 (Dengue 1 05K3903DK1) (GenBank accession number EU081242) was isolated during the 2005 dengue outbreak in Singapore. DENV2 (Dengue D2Y98P) (GenBank accession number JF327392) derives from a clinical strain isolated in Singapore in 1998 that had been exclusively passaged in C6/36 cells and plaque purified twice in BHK-21 cells. EHIE2862Y15 (GenBank accession number MK513444) was a DENV2 clinical strain isolated in Singapore in 2015. All DENV stocks were propagated in C6/36 cell line maintained in Leibovitz’s L-15 medium supplemented with 2% FBS as previously described (Ng et al., 2014). Harvested culture supernatants containing the virus particles were stored at −80 °C. Virus titers were determined by plaque assay in BHK-21 cells as described below.

### Virus quantification

Virus titer was quantified by plaque assay in BHK-21 cells as previously described (Ng *et* al., 2014). Briefly, 40,000 cells/well were seeded in 24-well plates (NUNC) one day before plaque assay. Cell monolayers were then infected with 10-fold serially diluted viral suspensions in RPMI-1640 medium supplemented with 2% FBS. After one hour incubation at 37 °C with CO_2_, overlay medium (RPMI-1640 medium containing 1% (w/v) carboxymethyl cellulose (CMC) and 2% FBS) was added to each well. After incubation for four (DENV2) or five (DENV1) days at 37 °C with CO_2_, cells were fixed with 4% paraformaldehyde (Sigma-Aldrich) and stained with 0.05% crystal violet (Sigma-Aldrich). Plaques were counted, adjusted by dilution and expressed as the number of plaque forming unit per milliliter (PFU/mL).

### Chimeric virus construction and production

Fragments from DENV2 (D2Y98P) and DENV1 (05K3903DK1) genomes were obtained by polymerase chain reaction (PCR) amplification or *de novo* synthesis. Primer sequences used for amplification are listed in Table S3. PCR was performed using Q5© Hot Start high-fidelity DNA polymerase (New England Biolabs) according to the manufacturer’s instructions. All PCR products were purified from agarose gel using QIAquick® gel extraction kit (Qiagen) following the manufacturer’s instructions. DENV1-NS1-D2Y98P and DENV1-prME-D2Y98P chimeric viruses were constructed using DNA assembly of PCR fragments by Gibson assembly technique as previously described (Siridechadilok et al., 2013). The pACYC177 plasmid containing CMV promoter, HDV ribozyme and SV40 PA was a kind gift from Prof Ooi Eng Eong, Duke-NUS. The vector and viral DNA fragments, were assembled together using Gibson Assembly® Master Mix (New England Biolabs) at 50 °C for two hours. The assembly mix (15 μL) was then directly transfected into BHK-21 cells using Lipofectamine-2000® transfection reagent (Thermo-Fisher) in Opti-MEM medium (Thermo-Fisher) as described in the manufacturer’s protocol. After incubation for four hours at 37 °C with CO_2_, the medium was changed to MEM medium (GIBCO) supplemented with 10% FBS. Culture supernatants containing the virus were collected daily from day one to day five post-transfection and the presence of virus was verified by plaque assay. The chimeric viruses were eventually propagated in C6/36 cell line for two passages.

### NS1 immunization and mouse infection

Five to six week-old A129 mice (129/Sv mice deficient in IFN-α/β receptors) were immunized intraperitoneally (ip) thrice (week 0, 2 and 6) with 20 μg of hexameric NS1 from DENV2 16681 strain (The Native Antigen Company) or OVA protein (InvivoGen) adjuvanted with 1 μg of monophosphoryl lipid A (MPLA) (InvivoGen) and 1:1 volume AddaVax (InvivoGen) as described previously (Beatty et al., 2015). Two weeks after the last administration, A129 mice were challenged with 10^6^ PFU DENV2 (D2Y98P) intravenously (iv) one day after iv administration of DENV1-immune serum to trigger ADE.

Following the same NS1(16681) immunization regimen in A129 mice, NS1 immune serum was collected two weeks after the third immunization and heat-inactivated at 56 °C for 30 minutes before storage at −80 °C. Antibody titers of immune serum were quantified by ELISA. The stored immune sera were subsequently used for passive transfer experiment into AG129 mice (129/Sv mice deficient in IFN-α/β and IFN-γ receptors) born to DENV1-immune mothers. One day post-administration of NS1 immune serum (150µL per mouse), these AG129 mice were subcutaneously (sc) challenged with 10^3^ PFU of DENV2 (D2Y98P).

Five to seven week-old IFNAR^-/-^ mice (C57BL/6 deficient in interferon-α/β receptors) were subjected to the same immunization regimen with NS1(16681) or OVA protein as described above, and were challenged sc with 10^5^ PFU DENV2 (D2Y98P or EHIE2862Y15) two weeks after the third immunization.

### Administration of NS1 protein

Five to six weeks old AG129 mice were infected sc with 10^3^ PFU sub-lethal dose of D2Y98P. Two or four days p.i., 10 mg/kg of purified NS1(D2Y98P) (The Native Antigen Company) or OVA protein (InvivoGen) was administered iv to the infected mice.

### Primary infection with chimeric viruses

Five to six weeks old AG129 mice were sc infected with 10^6^ PFU of parental (D2Y98P or DENV1) or chimeric (DENV1-NS1-D2Y98P or DENV1-prME-D2Y98P) viruses.

### Quantification of vascular leakage

Vascular leakage was assessed by Evans blue dye extravasation as previously described (Ng et al., 2014). Briefly, mice were administered intravenously with 0.5% (w/v) Evans blue dye (Sigma-Aldrich) in PBS adjusted to the weight of mouse (10 μL/gram body weight). Two hours after administration, mice were euthanized and extensively perfused with PBS. Organs (liver, small intestines, spleen and kidneys) were harvested and weighed. Evans blue dye was extracted from the organs by the addition of N,N-dimethylformamide (Sigma-Aldrich) (adjusted to 4 mL/g wet tissue) and incubated overnight at 37 °C. Extracted dye was read at 620 nm, and data were expressed as absolute absorbance.

### Histology analysis

Mice were sacrificed at the indicated time point and the liver was harvested and fixed immediately with 4% paraformaldehyde in PBS. Fixed tissues were processed for embedding, sectioning and staining with Haematoxylin and Eosin (H&E) (Department of Pathology, NUS). Slides were viewed under microscope (Leica) and images were captured.

### Enzyme-link immunosorbent assays

Levels of systemic IgG antibodies specific to NS1 protein were quantified via indirect enzyme-link immunosorbent assay (ELISA). Purified NS1 protein from D2Y98P or 16681 (10 ng/well) diluted in PBS was coated onto 96-well EIA plates (Corning costar) overnight at 4 °C. Plates were washed thrice with wash buffer (0.05% Tween 20 in PBS) and blocked with reagent diluent (2% BSA in wash buffer) for one hour at 37 °C. Serially diluted serum samples were added to the wells and incubated at 37 °C. Plates were washed thrice before the addition of horseradish peroxidase (HRP)-conjugated anti-mouse IgG (H+L) (Bio-rad, 170-6516) at 1:3,000; anti-mouse IgG1, IgG2a and IgG2b (abcam ab97240, ab97245, ab97250) at 1:10,000. Plates were incubated for one hour at 37 °C. After the final three washes, detection was performed by the addition of *o*-phenylene-diamine dihydrochloride substrate SigmaFast (Sigma-Aldrich) and incubated for 30 minutes at room temperature. The reaction was stopped upon the addition of 2 N H_2_SO_4_. Absorbance was read at 490 nm, and antibody titer was determined by non-linear regression as the reciprocal of the highest serum dilution with absorbance corresponding to three times the absorbance of blank wells.

Levels of systemic sNS1 in mice during the course of infection or NS1 in *in vitro* experiment were quantified via sandwich ELISA as described previously (Watanabe et al., 2012). Mouse anti-NS1 Mab62.1 (a kind gift from Prof Subhash Vasudevan, Duke-NUS and Prof Christiane Ruedl, Nanyang Technological University) (0.1 μg/well) diluted in PBS was coated onto 96-well EIA plates overnight at 4 °C. After washing and blocking as described above, diluted serum samples (from 1:100 to 1:2,500) were added to the wells and incubated for 2.5 hours at 37 °C. Standard curve was established by two-fold serial dilution of recombinant NS1 (D2Y98P or 16681) from 25 ng/mL to 0.39 ng/mL. Plates were washed five times before the addition of HRP-conjugated mouse anti-NS1 MAb56.2 (a kind gift from Prof Subhash Vasudevan, Duke-NUS and Prof Christiane Ruedl, Nanyang Technological University) (25 ng/well, made by conjugating HRP to MAb56.2 using NH_2_ peroxidase labelling kit (Abnova) and incubated for 1.5 hours at room temperature. After the final washes, detection was performed with the addition of tetramethylbenzidine (R&D Systems) for 30 minutes at room temperature. The reaction was stopped with 2 N H_2_SO_4_. Absorbance was read at 450 nm. The concentration of NS1 was calculated based on the standard curve and expressed as the concentration of NS1 in ng/mL.

Levels of circulating TNF-α in infected mice were measured using Mouse TNF-α Immunoassay Quantikine® ELISA kit (R&D Systems) according to the manufacturer’s instructions. Levels of heparan sulphate in circulation were quantified using Mouse Heparan Sulfate ELISA kit (G-Biosciences) according to the manufacturer’s instructions.

### Multiplex cytokine and soluble mediators detection

Levels of cytokines and soluble mediators present in plasma of infected mice were quantified using LEGENDplex™ Mouse Inflammatory Panel kit (Biolegend) according to the manufacturer’s protocol. Samples were run using Attune™ Nxt flow cytometer and analyzed using FlowJo software.

### *In vitro* DENV infection

Vero cells were infected at a multiplicity of infection (MOI) of 0.1 with D2Y98P, DENV1, DENV1-prME-D2Y98P and DENV1-NS1-D2Y98P. Plates were incubated at 37 °C for one hour with rocking every 15 minutes interval for viral adsorption. Each well was rinsed twice with PBS before addition of 200 μL of DMEM medium containing 2% FBS. The plates were incubated for four days and the culture supernatant was collected at indicated time points post-infection. Viral quantification was performed by plaque assay.

### Statistical analysis

Data analyses were performed using Graphpad Prism 6.0. Statistical comparison was conducted using non-parametric Mann-Whitney test or linear regression. Comparison of survival rates was performed using Log-rank (Mantel-Cox) test. Differences were considered significant (*) at *p* value <0.05.

## Author Contributions

PXL, DHRT, CLPHB, ETXT, JZHC, LCO, YLC, CH performed the experiments; PXL, DHRT, SA designed the experiments and wrote the manuscript; PXL, DHRT, CH, LCN, SA analysed the data.

## Acknowledgements

We thank Prof Subhash Vasudevan, Duke-NUS and Prof Christiane Ruedl, Nanyang Technological University for the kind gift of mouse monoclonal anti-NS1 antibodies for ELISA. We also thank Prof Ooi Eng Eong, Duke-NUS for sharing his pACYC177 plasmid construct for Gibson assembly work.

This work was funded by the Singapore Ministry of Health (CBRG13nv005 and MOHIAFCAT2/3/011 awarded to SA).

## Supplemental Information

**Fig. S1.**
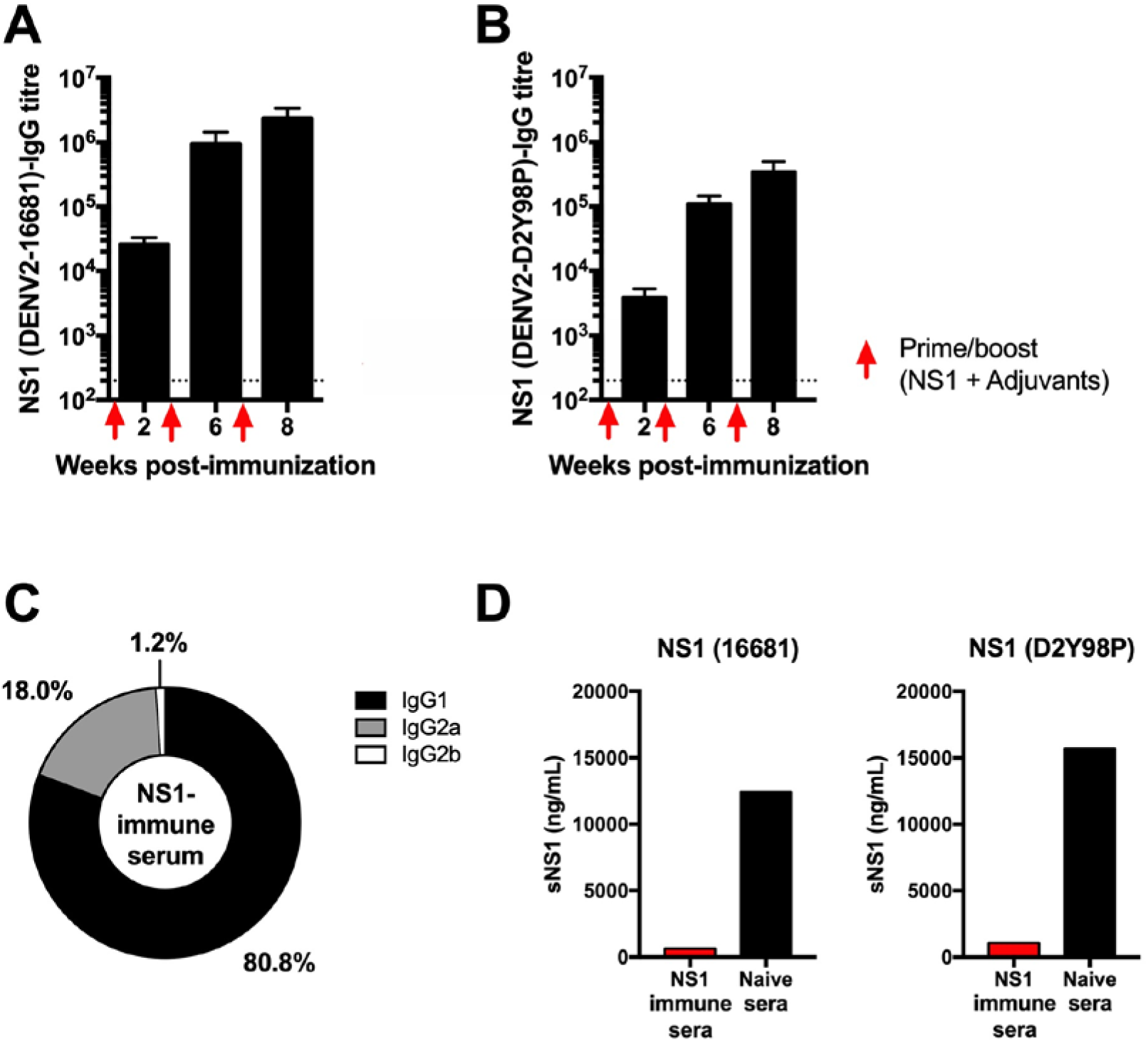
NS1-specific IgG titers in NS1 immune serum. A129 mice were ip immunized thrice with 20 μg NS1(16681) or OVA adjuvanted with MPLA and Addavax at week two, six and eight. NS1-specific IgG antibody titers were measured by ELISA using NS1(16681) (A) or NS1(D2Y98P) (B) as coating antigen. Titers were calculated as the reciprocal of the highest serum dilution with absorbance corresponding to three times the absorbance of blank wells. The dotted line indicates the limit of detection. (C) The percentages of specific IgG subclasses were determined. (D) 5x-diluted NS1 immune serum was co-incubated with 20µg/mL NS1(16681) or NS1(D2Y98) and the amount of free NS1 was measured by sandwich ELISA.

**Fig. S2.**
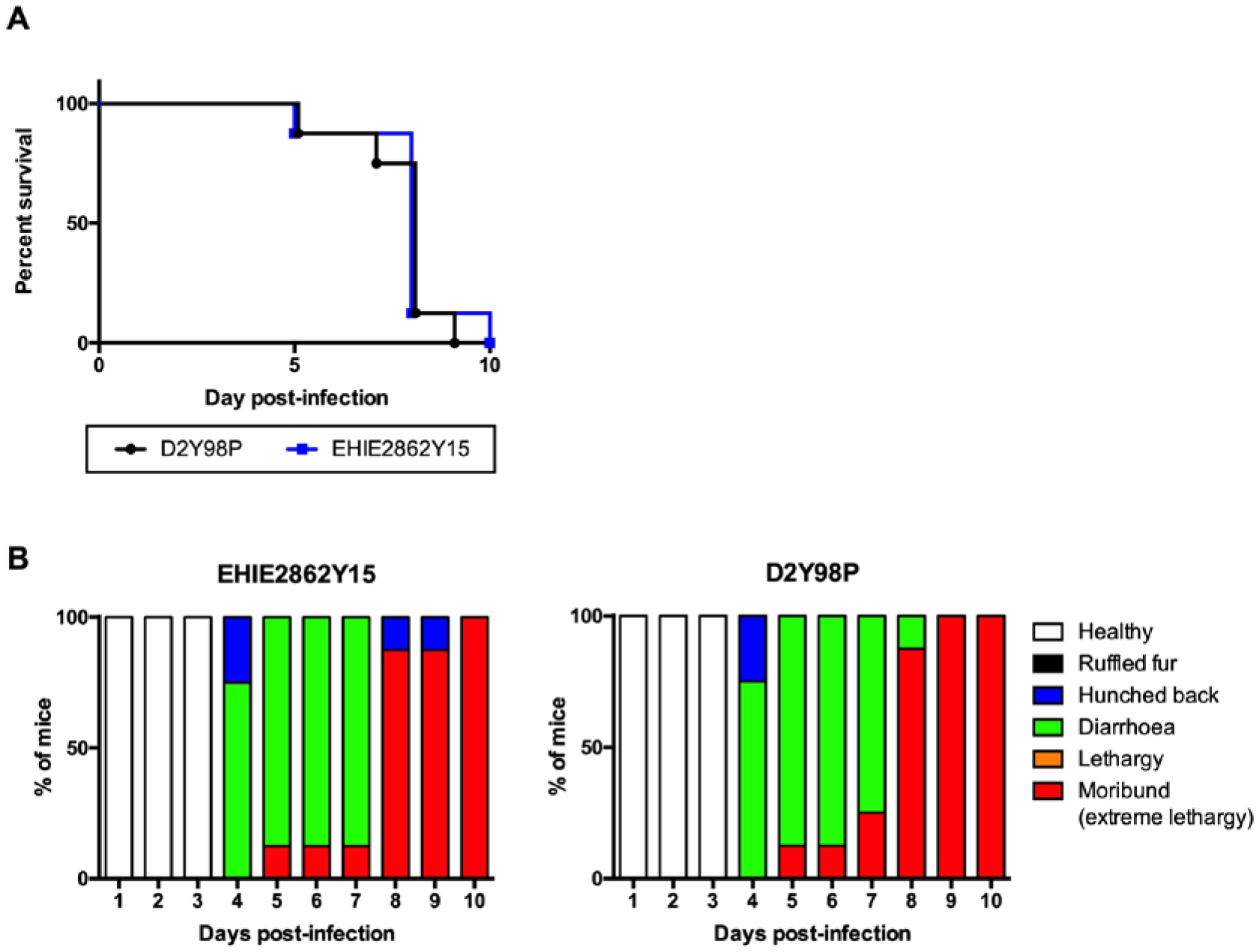
*In vivo* fitness of EHIE2862Y15 strain in AG129 mice. AG129 mice (n = 8) were sc challenged with 10^6^ PFU of EHIE2862Y15 or D2Y98P virus. Mice were monitored daily upon challenge. The Kaplan-Meier survival curve (A) and clinical scores (B) are shown.

**Fig. S3:**
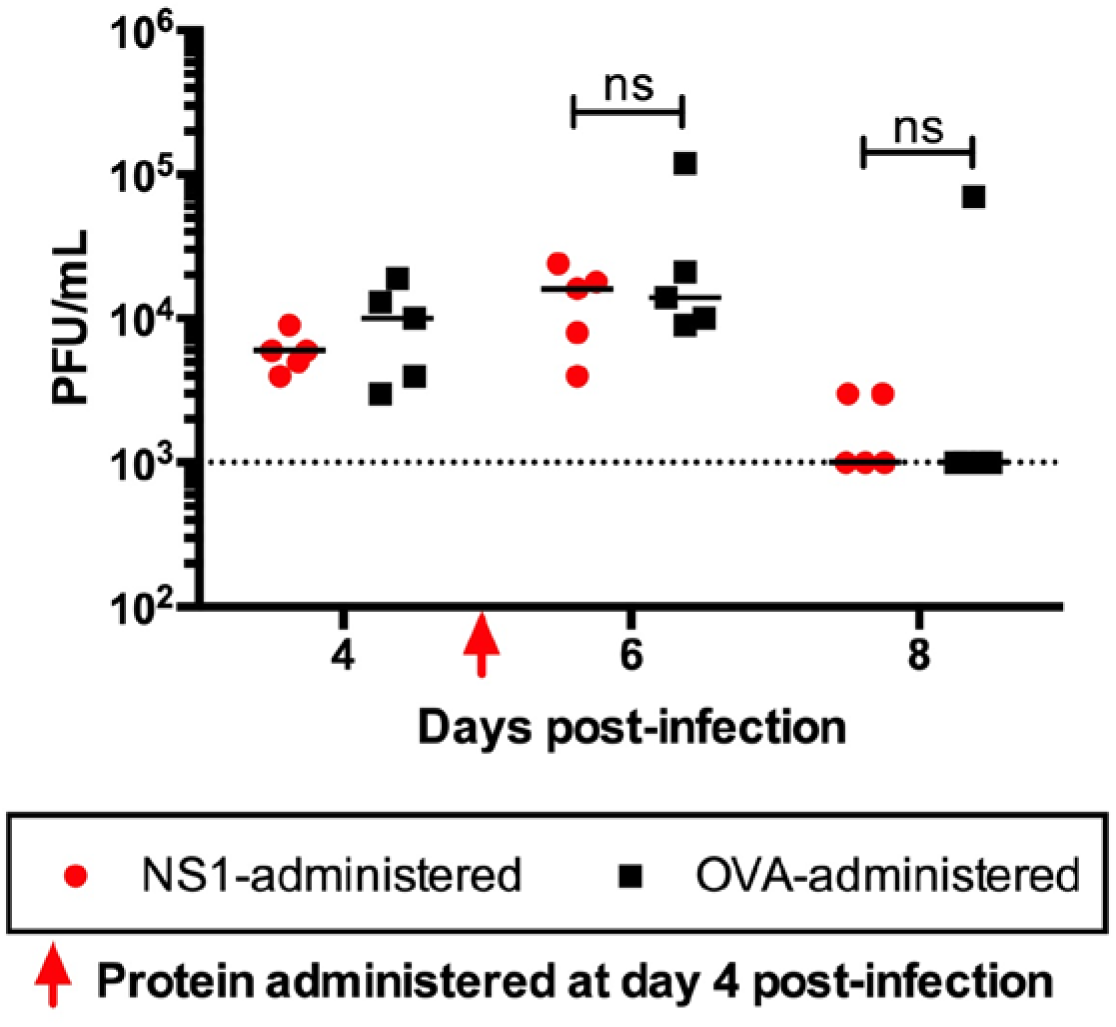
Viremia titers in D2Y98P-infected AG129 mice administered with NS1. AG129 mice were sc challenged with 10^3^ PFU D2Y98P. At day 4 p.i., D2Y98P-infected mice were iv administered with 10mg/kg NS1(16681) or OVA. Dotted line denotes the limit of detection. Data were analyzed by non-parametric Mann Whitney test. ns: not significant.

**Fig. S4.**
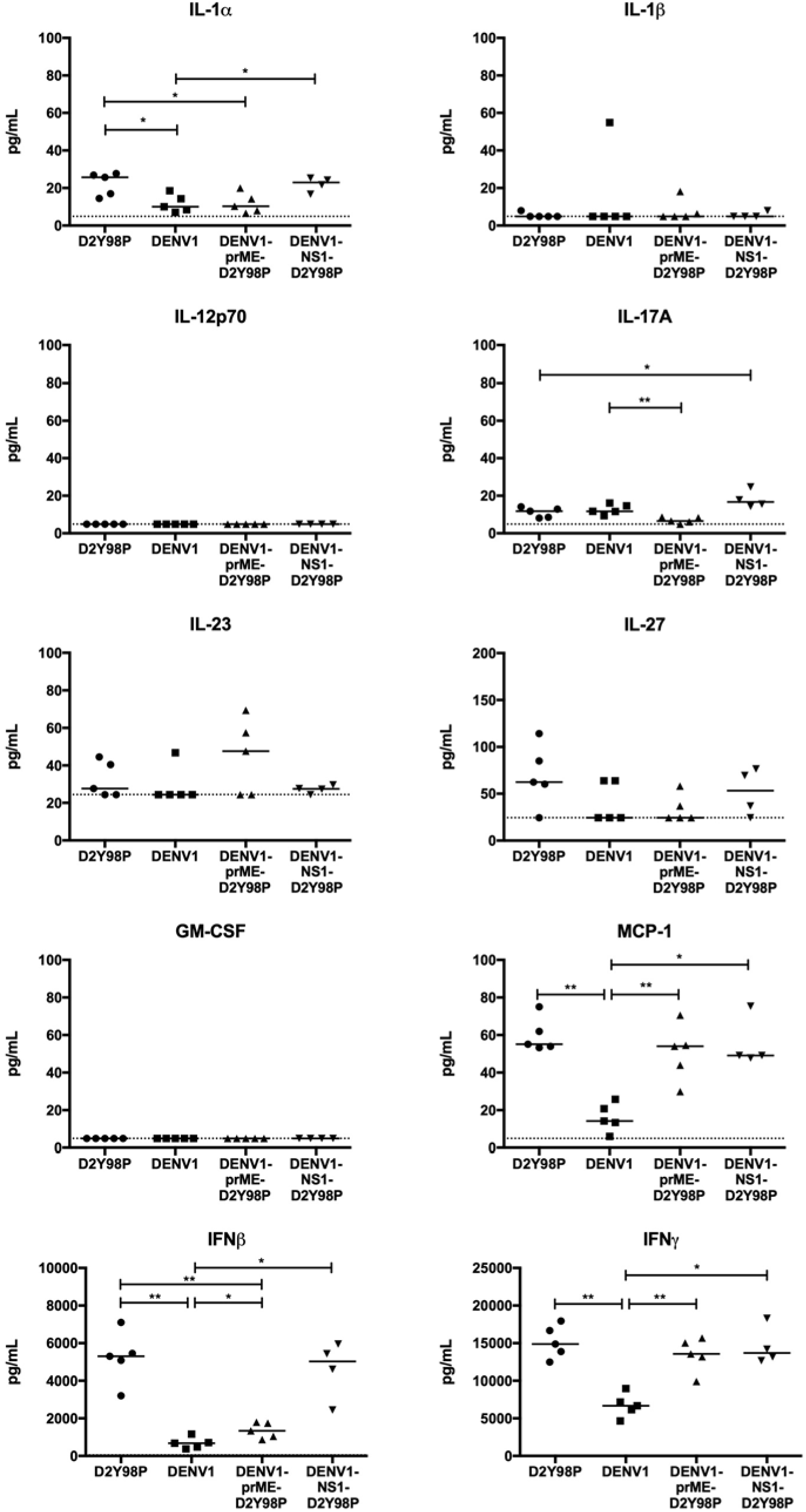
Cytokine profile in AG129 mice infected with chimeric and parental viruses. AG129 mice (n = 4-5) were sc infected with 10^6^ PFU of D2Y98P, DENV1, DENV1-prME-D2Y98P or DENV1-NS1-D2Y98P virus. At day 5 p.i., the systemic levels of various cytokines was measured. Data were analyzed by non-parametric Mann-Whitney test. *\*p*<0.05; *\*\*p*<0.05.

**Table S1.**
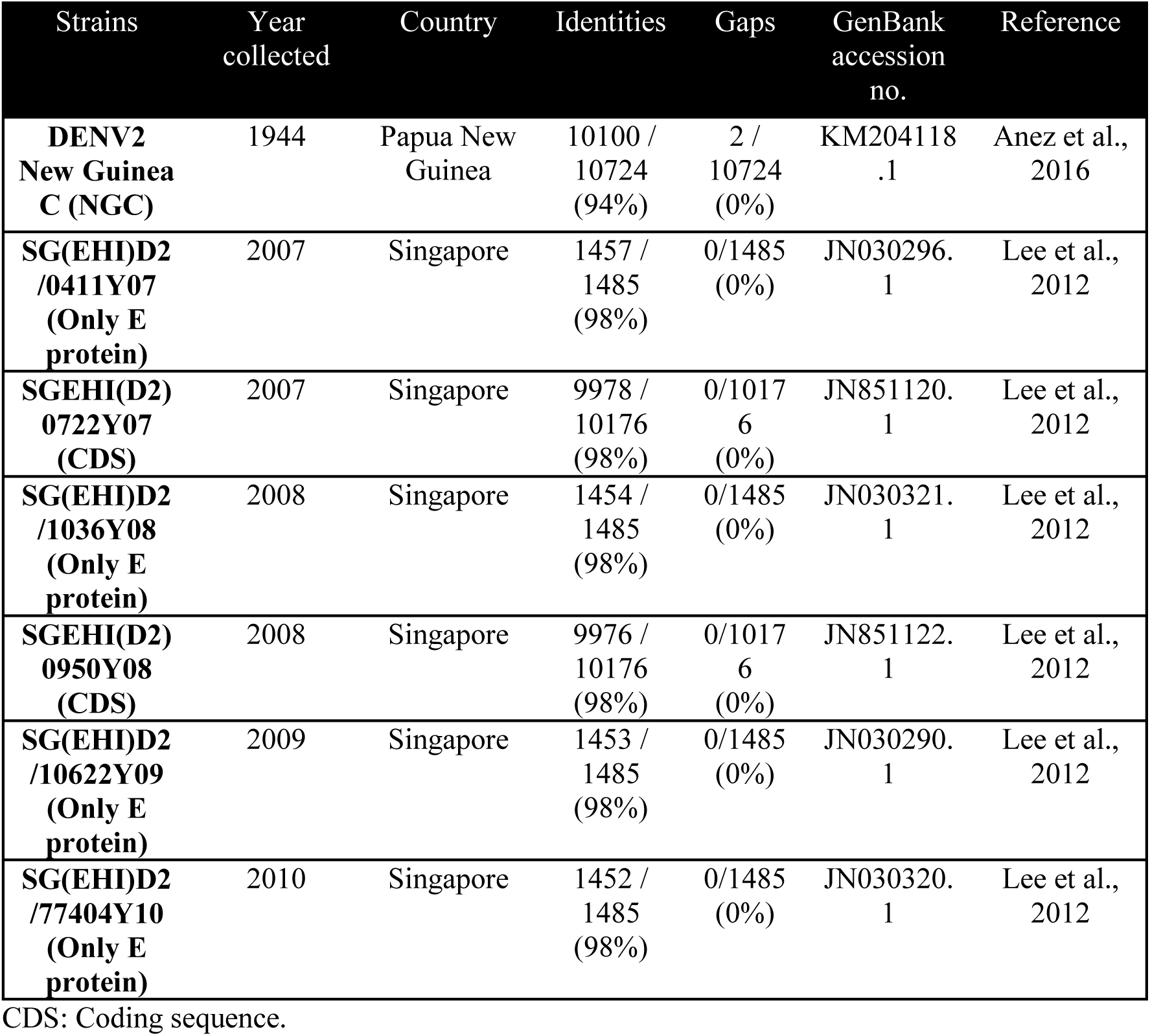
Comparison of D2Y98P genome against DENV2 NGC strain and Singapore DENV2 clinical isolates.

**Table S2.** Amino acid sequence comparison between EHIE2862Y15 and D2Y98P strains.

| **EHIE2862Y15**<br>**vs D2Y98P** | **Identity** | **Positive** | **DENV Protein** | **Differences** |
| --- | --- | --- | --- | --- |
| **Nucleotide**<br>**Level** | 10570/10723<br>(98.6%) |  |  |  |
| **Amino Acid**<br>**Level** | 3354/3368<br>(99.6%) | 3359/3368<br>(99.7%) | NS1 | T80S (+)<br>S178F |
|  |  |  | NS2A | A171T<br>T189A |
|  |  |  | NS2B | F21L<br>I59V (+) |
|  |  |  | NS3 | R15K (+) |
|  |  |  | NS4A | V89I (+) |
|  |  |  | NS4B | L52F |
|  |  |  | NS5 | V271T<br>T414N<br>S585P<br>A637V<br>N676S (+) |

**Table S3.**
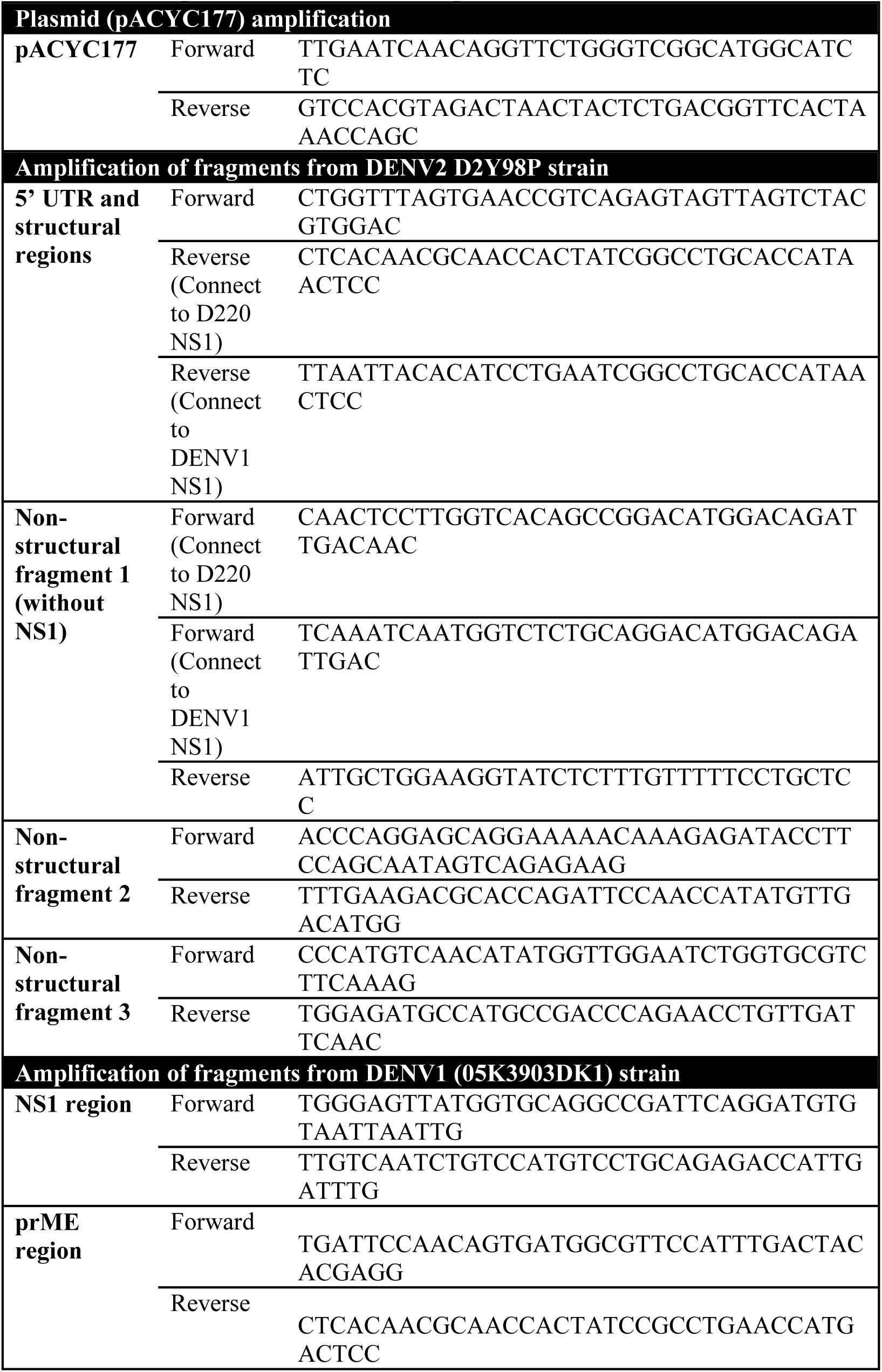
List of primers used for PCR amplification.

